# Disgust propensity, not disgust sensitivity, shapes the reactivity of a subjective disgust circuit in humans

**DOI:** 10.1101/2025.08.27.672493

**Authors:** Xianyang Gan, Zihao Zheng, Ran Zhang, Feng Zhou, Ting Xu, Nan Qiu, Junjie Wang, Heng Jiang, Shan Gao, Yu Wu, Benjamin Klugah-Brown, Dezhong Yao, Benjamin Becker

## Abstract

Disgust constitutes an evolutionary adaptive defensive-avoidance response, yet humans vary markedly in their dispositional tendency to experience disgust (disgust propensity) and in their negative appraisal of such experience (disgust sensitivity). Conceptual frameworks and neuroimaging studies suggest that these traits may differentially modulate neural responses to disgust-eliciting stimuli; however, methodological constraints have left their precise roles unresolved. Our comparably large fMRI study (n = 142) therefore aimed to systematically determine how trait disgust modulates neural responses to carefully selected and validated disgust-specific visual stimuli across varying levels of subjective disgust experience. The whole-brain voxel-wise regression analyses revealed a neural dissociation between the two disgust traits, with disgust propensity, but not disgust sensitivity, modulating disgust-related neural activity in the anterior, middle, and posterior insula, as well as the caudate, putamen, thalamus, hippocampus, and parahippocampal gyrus. Mediation and network-level analyses further supported this dissociation by showing that disgust propensity shapes disgust experience via insula – striatal – hippocampal pathways. Together, these findings provide evidence for a neurofunctional dissociation of disgust propensity and sensitivity and elucidate how trait disgust shapes subjective experiences. They further suggest that disgust propensity and the identified systems may represent promising targets for the regulation of disgust-related pathology.

## 1. Introduction

Disgust has been conceptualized as an evolutionary adaptive defensive-avoidance response, which may have initially evolved in mammalian rejection of potentially poisonous food [1-4]. Consistent with its proposed evolutionary defensive-avoidance function, the feeling of disgust is primarily elicited by poisonous as well as contaminated and contagious stimuli, indicating a broader function in pathogen avoidance [5-8].

Over the past three decades, numerous studies have used functional magnetic resonance imaging (fMRI) to examine the neural substrates of disgust in humans. The majority of studies in this field typically present participants with disgust-eliciting stimuli from different sensory modalities and average the participants’ neural responses on the group level. These studies enable us to identify the core brain systems involved in disgust processing, with recent large-scale meta-analytic syntheses showing that disgust engages the defensive-avoidance circuit, including the insula, amygdala (extending into the parahippocampal gyrus), basal ganglia, as well as prefrontal, parietal, and temporo-occipital regions [6,9].

However, these studies did not take into account individual differences in trait disgust, such as disgust propensity – the general tendency to respond with disgust in a certain situation – or disgust sensitivity, referring to how unpleasant the experience of disgust is appraised [10,11]. Individuals with greater disgust propensity are more likely to perceive stimuli as disgust-eliciting [11], which may result in stronger activation of disgust-related brain regions compared to individuals with lower levels of disgust propensity [12]. Elevated levels of disgust propensity have been linked to behavioral avoidance [11,13-15], and together, these factors have been implicated in the etiology of various psychopathological conditions (e.g., eating disorders and, in particular, different forms of anxiety disorders; see [16-21]). However, in the development of psychopathology, what matters may not only involve disgust propensity but also disgust sensitivity [10,11]. Notably, both types of trait disgust have been shown to be differentially associated with symptoms of anxiety disorders [10,11,15,22-24]. Taken together, these findings highlight the importance of examining how trait-level individual differences in disgust propensity and sensitivity shape the neural substrates of disgust.

In response to this need, another line of research has utilized correlational approaches (e.g., regression analyses) to investigate how individual differences in trait disgust modulate neural responses during disgust induction. However, findings remained highly inconsistent. The insula has been widely recognized as a key region associated with disgust propensity [7,16]. Some studies observed an association between trait disgust propensity and activation of the insula to disgust-eliciting stimuli [12,25], while other studies failed to observe such results [26-28]. Furthermore, three early studies compared the modulatory roles of both disgust traits (propensity and sensitivity) and yielded conflicting findings. Two of them revealed an association between disgust sensitivity, rather than disgust propensity, and insula activity in response to disgust stimuli [29,30], whereas the third study found the opposite result - that disgust propensity, not sensitivity, modulated insula activity [31]. These discrepancies may be due to experimental and technical limitations of the previous studies. First, none of the studies collected participants’ momentary subjective disgust experience evoked by each disgust stimulus while undergoing fMRI (also termed as ‘state disgust’), but instead employed the categorical approach of comparing a priori assigned high disgust versus neutral stimuli. The neglect of incorporating individual differences in state disgust into the statistical model may inherently capture neural activity related to salience, arousal, or autonomous reactivity, thereby obscuring the correspondence between subjective feelings and their brain representation [5,32-35]. Second, the majority of visual stimuli used to induce disgust experience in most of the studies were selected from standardized affective picture databases (e.g., International Affective Picture System, IAPS [36]) that were not specifically designed to induce disgust and may also induce high levels of general negative affect [25], a response that is distinguishable from subjective disgust experience on the neural level [5,37]. Finally, all studies - except for Mataix-Cols et al. (2008) [25] - examining associations between trait disgust and neural reactivity focused on a priori defined brain regions of interest (ROI) analyses and did not utilize a hypothesis-free and more conservative voxel-wise whole-brain correlation approach.

The present study thus seeks to determine how individual differences in both disgust propensity and disgust sensitivity influence neural responses to a diverse set of validated disgust-specific affective pictures (from the DIsgust-RelaTed-Images (DIRTI) disgust images database [38]; for details, see Methods) in a large sample of healthy individuals (n = 142). Importantly, the included pictures span a wide range of disgust intensities and are expected to evoke varying levels of subjective disgust experience, assessed online during fMRI (details see [5]). Briefly, we utilized the behavioral and neural data to determine the modulatory role of trait disgust on neural responses under varying levels of state disgust, as captured by multiple contrast-based comparisons (e.g., high disgust > neutral, high disgust > moderate disgust), using a whole-brain voxel regression approach. We subsequently employed mediation and network-level analyses to link trait variations with subjective experience and neural responses. Specifically, we conducted a series of mediation analyses to further examine the relationship between state disgust, brain activation, and trait disgust. In addition, network-level analyses were performed by integrating meta-analytic connectivity mapping (MACM) and resting-state functional connectivity (RSFC) to delineate the connectivity profiles of brain regions identified through the core regression analyses. Overall, these systematic analyses enable us to achieve a more comprehensive characterization of how trait disgust shapes neural responses to disgust-inducing stimuli.

## 2. Methods

### (a) Participants

A total of 146 healthy, right-handed participants with normal or corrected-to-normal vision were initially recruited. Exclusion criteria included MRI contraindications, neurological or psychiatric disorders, medication or substance use. Four participants were excluded for head motion >3 mm, resulting in a final sample of 142 (75 females; mean ± s.d. age 22.23 ± 2.60 years). The study was part of a larger project (details see [5]) and was approved by the Ethics Board of the University of Electronic Science and Technology of China. Written informed consent was obtained from each participant before the experiment.

### (b) Self-report trait disgust questionnaire

We chose the Disgust Propensity and Sensitivity Scale-Revised (DPSS-R [10]), instead of content-dependent trait disgust measures such as the Disgust Scale [39] or the Questionnaire for the Assessment of Disgust Proneness [40], given that the DPSS does not contain specific disgust elicitors, which is vital to prevent spurious associations between trait disgust scores and actual state disgust responding during the experiment owing to content overlap of stimuli [29,30]. Furthermore, it is the first scale that meaningfully differentiates between disgust propensity and disgust sensitivity [10]. The DPSS-R contains 16 items constituting two validated trait disgust subscales: an 8-item disgust propensity subscale and an 8-item disgust sensitivity subscale. The translation of the DPSS-R into Chinese adhered to a strict forward and backward translation procedure by bilingual English Professors from the school of foreign languages and psychological professionals. The final iteration yielded a scale assumed to accurately capture the context-adjusted English items of DPSS-R.

Prior to scanning, participants completed the Chinese DPSS-R (available via email to the corresponding author), rating each statement on a 5-point Likert scale (1 = never, 5 = always). Example items include ‘Disgusting things make my stomach turn’ (propensity) and ‘I worry that I might swallow a disgusting thing’ (sensitivity). In our sample, the Cronbach’s α coefficient for the disgust propensity subscale was 0.80, and the Cronbach’s α coefficient for the disgust sensitivity subscale was 0.77, both of which were highly similar to the original English version of DPSS-R (i.e., Cronbach’s α = 0.78 for the disgust propensity subscale, and Cronbach’s α = 0.77 for the disgust sensitivity subscale; for more details see [10]), indicating a reliable Chinese version of DPSS-R.

### (c) fMRI task

The present study encompassed two disgust fMRI experiments with (partly) different stimuli that were originally designed as a discovery-replication design (see [5]). The first experiment included 80 pictures that could elicit varying levels of subjective disgust experience from the DIRTI database [38] (67 pictures), the IAPS database [36] (8 pictures), and the Nencki Affective Picture System [41] (5 pictures), distributed over four runs (with 20 stimuli in each run). To prevent category-driven specificity in disgust, we selected a balanced sample from animals, scenes, and humans. Each picture was presented for 6 s, participants were instructed to look at the pictures and naturally experience the evoked emotion. After the presentation of each picture, a 2 s fixation cross was shown, followed by a 4 s response window during which participants rated their subjective disgust on a 5-point Likert scale (1 = neutral/slightest disgust; 5 = most intense disgust). Trials were separated by a jittered fixation cross epoch between 5 and 7 s. E-Prime 2.0 was used to present the stimuli. The second experiment contained 60 pictures, all drawn from the DIRTI database [38], distributed over two runs (with 30 stimuli per run). Each picture was displayed on the screen for 6 s, during which participants were required to pay attention to the pictures and naturally experience the elicited emotion. Then, a fixation cross jittered between 3 and 5 s was shown on the screen. Next, participants had 4 s to report the level of disgust they experienced for the stimuli on a 5-point Likert scale (1 = neutral/slightest disgust; 5 = most intense disgust). Trials were separated by a jittered fixation cross epoch between 5 and 7 s. E-Prime 3.0 was used to present the stimuli. Among the 142 participants (after excluding participants with excessive head movement), 108 finished the first experiment, while the remaining 34 participants completed the second experiment. Moreover, 133 out of 142 participants reported all five levels of subjective state disgust, whereas the remaining 9 reported ratings ranging only from level 1 to level 4. Before each experiment, participants performed a practice task outside the scanner to familiarize themselves with the experimental procedure. Both experiments reliably elicited the full range of subjective disgust ratings [5] (see Results for more information).

See Supplementary Methods for the details of MRI data acquisition.

### (d) fMRI data preprocessing

SPM12 (Statistical Parametric Mapping, https://www.fil.ion.ucl.ac.uk/spm/software/spm12/) was used to preprocess and analyze the fMRI data. The initial five volumes of fMRI data from each run were discarded to allow for T1 equilibration. Before preprocessing, we identified global motion outliers within each run based on either of the following criteria: (1) signal intensity larger than three s.d. from the global mean or (2) signal intensity and Mahalanobis distances larger than ten mean absolute deviations (https://github.com/canlab/CanlabCore). Each timepoint identified as outliers was subsequently incorporated into the first-level model as a separate nuisance regressor.

The preprocessing pipeline encompassed several key steps, including slice timing correction to account for temporal discrepancies across slices, realignment to correct for head motion, and unwarping to compensate for distortions caused by magnetic field inhomogeneities. Subsequently, high-resolution anatomical images underwent tissue segmentation into gray matter, white matter, cerebrospinal fluid, bone, fat, and air. The skull-stripped and bias-corrected structural image was then co-registered to the functional data, followed by normalization of the functional images to the standard Montreal Neurological Institute (MNI) space, resampled to an isotropic voxel size of 2 × 2 × 2 mm^3^. Finally, spatial smoothing was applied using an 8-mm full-width at half-maximum Gaussian kernel to enhance the signal-to-noise ratio and reduce spatial variability.

### (e) First-level fMRI analysis

Following preprocessing, first-level (subject-specific) analysis was conducted using the general linear model (GLM) framework [42]. Specifically, we constructed a GLM for each participant, which incorporated five or four regressors time-logged to the presentations of pictures for each rating level (i.e., 1-5 for those whose ratings covered from 1 to 5, or 1-4 for those who did not report rating 5) and one boxcar regressor aligned with the timing of rating period, thus allowing for the separate modeling of neural activity in response to each disgust level and for the modeling of effects associated with motor processes, respectively. The fixation-cross epoch served as an implicit baseline. Task-related regressors were convolved with the canonical hemodynamic response function, and for each voxel a high-pass filter with a cutoff of 128 s was implemented to remove low-frequency signal drifts. The four (for the first experiment) or two (for the second experiment) fMRI task runs were concatenated for each participant using the SPM function spm_fmri_concatenate.m. Nuisance regressors comprised run-specific intercepts, 24 motion parameters (six realignment parameters, their squares, derivatives, and squared derivatives per run), and vectors for outlier timepoints. Conditions of interest included high disgust (rating 4 and rating 5 for those whose subjective disgust ratings covered from 1 to 5, or rating 4 for those whose subjective disgust ratings only covered from 1 to 4), moderate disgust (rating 2 and rating 3), and neutral (rating 1). Contrast images between conditions (i.e., high disgust > neutral, high disgust > moderate disgust, and moderate disgust > neutral) used for the following regression analyses were estimated for each participant at each voxel, yielding statistical parametric maps that reflect the differential neural responses between conditions [42].

### (f) fMRI regression analysis

We employed a series of regression analyses to test whether variations in neural responses for each contrast were modulated by disgust propensity as well as disgust sensitivity. The resulting statistical maps were thresholded at *P* < 0.05 cluster-level family-wise error (FWE) correction, with cluster-forming voxel-level threshold at *P* < 0.001 (a minimum cluster size of 30 contiguous voxels was set to maintain spatial coherence), thereby obtaining regions significantly correlated with trait disgust scores.

### (g) Mediation analysis

To explore the relationship between state disgust, brain activation, and trait disgust, multiple mediation analyses were conducted via the SPSS Process v3.5. Here, state disgust was operationalized by the sum of participants’ self-reported subjective disgust ratings for each picture, brain activation was operationalized by the mean activation within each significant cluster derived from the above regression analyses (note that we only focused on those clusters that survived the stringent threshold, i.e., cluster-level FWE *P* < 0.05 correction and voxel-level *P* < 0.001; details see Results), trait disgust propensity was the sum of participants’ ratings for the disgust propensity subscale of the Chinese version of DPSS-R, and trait disgust sensitivity was the sum of participants’ ratings for the disgust sensitivity subscale of the Chinese version of DPSS-R. See Supplementary Methods for the statistical rationale of the mediation analysis. In this study, for each cluster we test whether disgust propensity (as well as disgust sensitivity) mediated the relationship between state disgust and the mean activation within this cluster. To evaluate the statistical significance of mediation effects, bootstrap tests with 10000 iterations were employed. If the bootstrapped 99% confidence interval does not include zero, the effect will be considered significant (*P* < 0.001). Given that the two disgust-induction fMRI tasks included differing numbers of pictures (that is, 80 in the first task and 60 in the second task), directly combining the raw total scores of subjective disgust ratings across these tasks would be biased. Therefore, total scores were normalized within each task before aggregation to yield a standardized measure of state disgust.

### (h) Task-based connectivity: MACM analyses

To investigate the meta-analytic co-activation patterns of each significant region obtained from the above regression analyses (details see Results), MACM analyses were conducted using the peak coordinates of each cluster identified in the regression analyses with the stringent threshold (cluster-level FWE *P* < 0.05 correction and voxel-level *P* < 0.001; details see Results) as ROIs (with sphere radius = 6 mm) based on the BrainMap functional database. Briefly, MACM examines whole-brain functional connectivity profiles by identifying the above-chance covariance between two or more brain regions [6,43]. This approach provides a measure of task-based functional connectivity by leveraging task-related neuroimaging data [44].

We performed a series of MACM analyses utilizing Sleuth software (v.3.0.4) from the BrainMap platform (https://brainmap.org/sleuth/). The search terms in the Sleuth were specified as follows: (1) “Locations: MNI Image” to upload the respective spherical ROI in MNI space, (2) “Experiments: Activations, Activations only” to restrict results to activation data, (3) “Experiments: Context, Normal Mapping” to focus on normative tasks, (4) “Subjects: Diagnosis, Normals” to include only healthy participants. This enabled the detection of above-chance co-activation patterns within a given seed ROI across a wide range of neuroimaging experiments from the functional database. After each respective ROI search, eligible activation coordinates from experiments were converted into MNI space and exported as a text file. The MACM analyses were based on the two datasets for each of the ROIs. Specifically, the dataset with the right anterior insula as ROI included 42 experiments, 801 foci, and 718 participants; and the dataset with the left thalamus as ROI encompassed 17 experiments, 229 foci, and 288 participants. To determine areas of convergence of co-activation with each ROI, these new datasets were analyzed separately using the activation likelihood estimation algorithm embedded in GingerALE v.3.0.2 (available via http://www.brainmap.org/ale/) with the following statistical criteria: voxel-level of *P* < 0.001 (uncorrected), cluster-level of *P* < 0.05 corrected, and 5000 permutations. See Supplementary Methods for the methodological overview of the activation likelihood estimation.

### (i) Task-free connectivity: RSFC analyses

To characterize intrinsic functional connectivity patterns of the regions identified through the above regression analyses (details see Results), RSFC analyses were performed on the original dataset, using 6 mm spherical ROIs centered on the peak coordinates of each significant cluster that survived the stringent threshold. Specifically, the original dataset used for the RSFC was from a separate cohort, consisting of 100 healthy subjects drawn from the Open Access Series of Imaging Studies (OASIS)-3 dataset (https://central.xnat.org). See Supplementary Methods for the data acquisition parameters and preprocessing procedures of this dataset.

Using the Data Processing Assistant for Resting-State fMRI (DPARSF, v.5.0_201001) (http://rfmri.org/DPARSF) toolbox, the preprocessed time series from each seed ROI was correlated with the time series of all other brain voxels. The resulting voxel-wise correlation coefficients were transformed into Fisher’s z-scores and tested for consistency across participants (thresholded at *P* < 0.05 cluster-level FWE correction, with cluster-forming voxel-level threshold at *P* < 0.001).

### (j) Consensus connectivity map

To identify overlapping connectivity patterns, MACM and RSFC results were integrated for each seed ROI using conjunction analyses based on the minimum statistic approach [45]. The resultant consensus connectivity maps identified brain regions showing consistent interactions with each seed across both task-based and task-independent functional connectivity profiles [9,46,47].

### (k) Spatial similarity between respective maps and thalamus subregions

To pinpoint the contributions of different thalamus subregions to the regression map as well as to the consensus connectivity maps (details see Results), we employed a sensitive approach that illustrates spatial similarity between respective maps and eight thalamus subregions derived from the Human Brainnetome Atlas [48]. Consistent with recent studies [5,37,49], spatial similarity was computed as cosine similarity between each thalamus subregion and the corresponding map obtained from regression and connectivity analyses.

## 3. Results

### (a) Self-report data

During fMRI, state disgust was evoked using validated disgust-specific affective pictures (the DIRTI database [38]). These pictures successfully elicited sufficient levels of subjective state disgust across the most common disgust-related categories (animal, human, and scene; Fig. 1), with over 11% of stimuli in each category inducing the highest level of state disgust (rating 5).

**Fig. 1.**
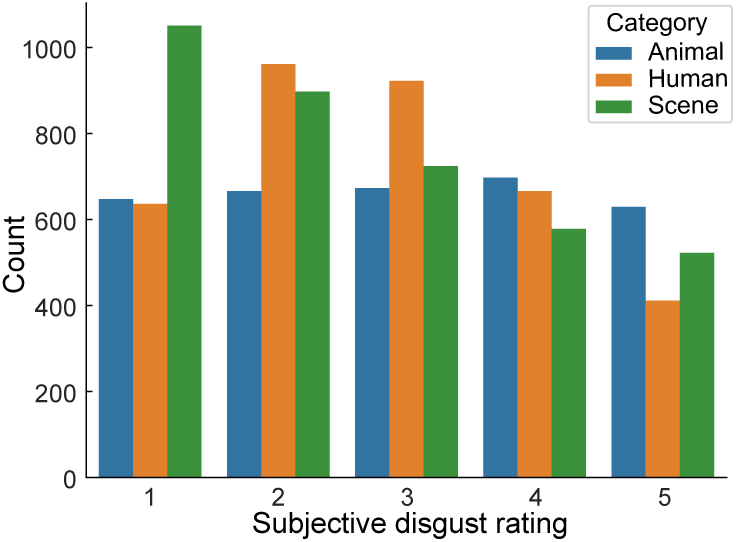
The distribution of state disgust ratings for each disgust category (animal, human, and scene).

Disgust propensity and disgust sensitivity were highly correlated (*r* = 0.71, *P* < 0.001), consistent with the original English version of DPSS (*r* = 0.59, *P* < 0.01; details see [10]). Moreover, both disgust propensity (*r* = 0.28, *P* < 0.001) and disgust sensitivity (*r* = 0.33, *P* < 0.001) were positively correlated with the experienced state disgust elicited by disgust stimuli.

Considering that prior research showed sex differences in disgust propensity [28], an independent samples *t* test was performed to compare disgust propensity between females and males. However, there were no significant sex differences in disgust propensity (female: 24.20 ± 4.43; male: 23.73 ± 5.08; *t*_(140)_ = 0.59, *P* = 0.56). Furthermore, we did not find sex differences in either disgust sensitivity (female: 18.60 ± 5.08; male: 19.05 ± 5.35; *t*_(140)_ = 0.51, *P* = 0.61) or the total scale score (female: 42.80 ± 8.65; male: 42.78 ± 9.82; *t*_(140)_ = 0.02, *P* = 0.99). In addition, females and males did not differ in state disgust (female: −0.11 ± 0.96 after normalization; male: 0.13 ± 1.02 after normalization; *t*_(140)_ = 1.46, *P* = 0.15).

### (b) Regression results

Whole-brain regression analysis showed that activation to high disgust stimuli (high disgust > neutral) was significantly modulated by participants’ disgust propensity scores in a cluster located in the right anterior insula, encompassing regions of the anterior, mid- and posterior insular cortex, caudate, and putamen as well as a cluster located in the left thalamus, encompassing regions of the thalamus, hippocampus, caudate, and parahippocampal gyrus (Table 1 and Fig. 2). In each of these regions, higher trait disgust propensity was associated with increased neural activation in response to high disgust stimuli. We did not find significant negative associations between participants’ disgust propensity scores and brain activation. Including sex as a covariate in the regression model did not qualitatively change the results (details see Table S1 and Fig. S1).

**Table 1.**
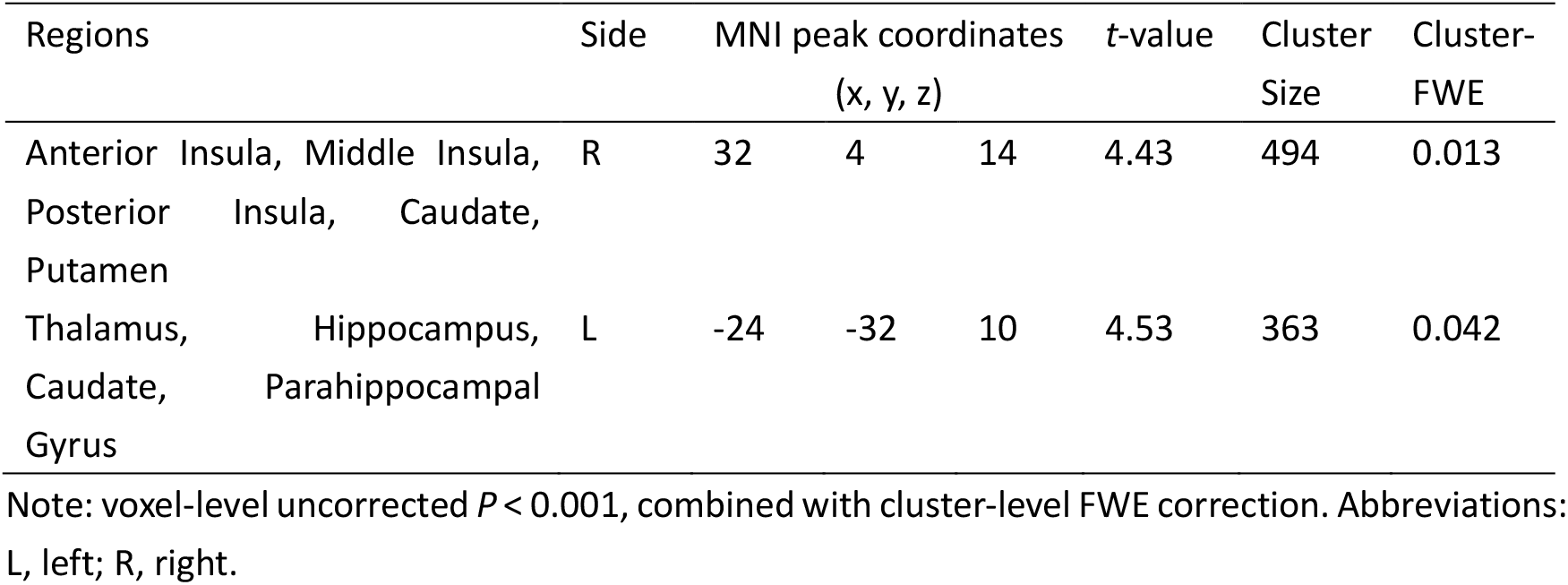
Brain areas where activation to viewing high disgust stimuli (high disgust > neutral) was significantly associated with participants’ scores on the disgust propensity subscale

**Fig. 2.**
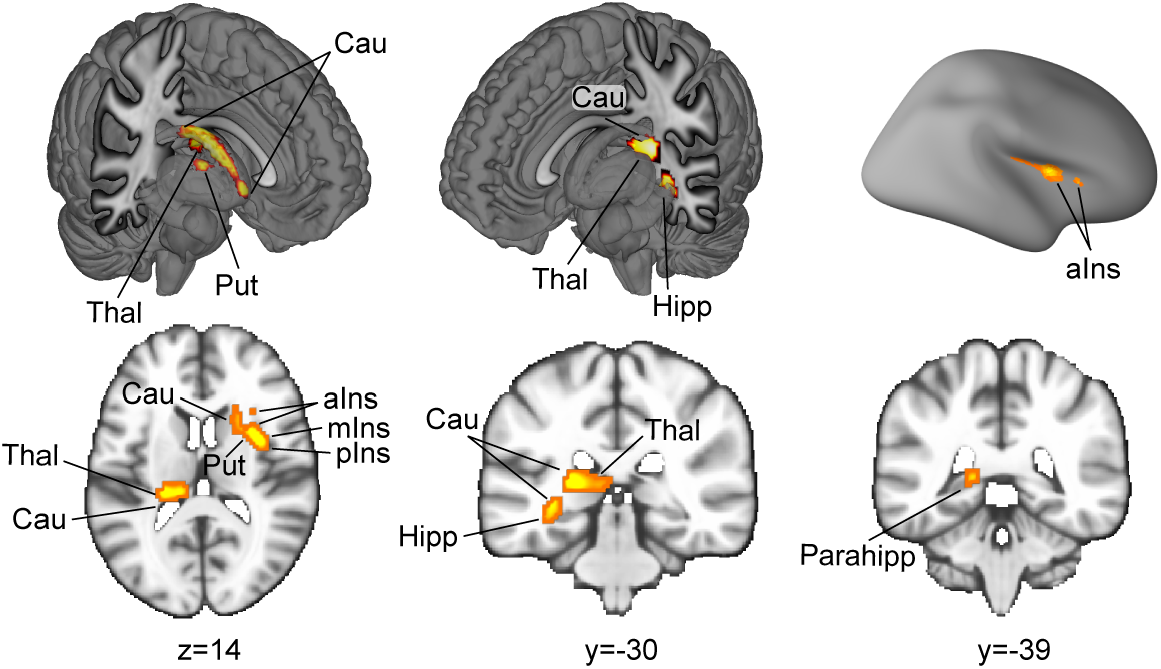
Positive association between disgust propensity scores and brain activation while viewing high disgust stimuli (high disgust > neutral). Note: activated voxel group shown at a statistical threshold of voxel-level *P* < 0.001 and cluster-level FWE *P* < 0.05 correction. Abbreviations: aIns, anterior insula; Cau, caudate; Hipp, hippocampus; mIns, middle insula; Parahipp, parahippocampal gyrus; pIns, posterior insula; Put, putamen; Thal, thalamus.

For the contrasts high disgust > moderate disgust and moderate disgust > neutral, no significant clusters emerged at cluster-level FWE *P* < 0.05 correction (voxel-level *P* < 0.001). Given this, we reported the regression results at the uncorrected voxel-level threshold (*P* < 0.001) to observe some trend. As for the condition of high disgust > moderate disgust, consistent with the findings reported above, activation in the right anterior insula and left thalamus clusters was again observed. Increased activation was also found in bilateral occipital cortices and left inferior/middle frontal gyrus (Supplementary Results, Table S2, and Fig. S2). For the condition of moderate disgust > neutral, significant modulation by participants’ disgust propensity scores was observed in a set of frontal and motor regions (Supplementary Results, Table S3, and Fig. S3). It should be noted that these results are suggestive due to the lenient threshold. Incorporating sex as a covariate did not qualitatively alter the above results.

However, regression analyses across the high disgust > neutral, high disgust > moderate disgust, and moderate disgust > neutral contrasts failed to identify any brain regions whose activation was significantly modulated by participants’ disgust sensitivity scores. These findings remained unchanged when sex was included as a covariate.

### (c) Mediation results

Regression analysis identified two clusters (right anterior insula and left thalamus; high disgust > neutral, cluster-level FWE *P* < 0.05, voxel-level *P* < 0.001, Table 1) with activation significantly associated with disgust propensity. Next, we performed a series of mediation analyses to test whether trait disgust mediated the relationship between state disgust and the mean activation in each cluster.

We first tested whether disgust propensity (*M*) could mediate the relationship between state disgust (*X*) and the mean activation of the anterior insula cluster (*Y*). As shown in Fig. 3a, state disgust was positively correlated with disgust propensity (path a; β = 1.34, SE = 0.39, 99% CI = [0.33, 2.35], *P* < 0.001), disgust propensity was positively associated with the mean activation of the anterior insula cluster independently of state disgust (path b; β = 0.03, SE = 0.01, 99% CI = [0.01, 0.04], *P* < 0.001) and there was a significant mediation effect (path a×b; β = 0.04, SE = 0.01, 99% CI = [0.008, 0.08], *P* < 0.001). Furthermore, the total effect (path c; β = −0.002, SE = 0.03, 99% CI = [-0.07, 0.07], *P* = 0.93) and the non-mediated effect (path c’; β = −0.04, SE = 0.03, 99% CI = [-0.11, 0.03], *P* = 0.13) were not significant. These results indicate that disgust propensity plays a full mediation role in the effect of state disgust on the mean activation of the anterior insula cluster. We next replaced disgust propensity with disgust sensitivity and ran a second mediation analysis. However, the results revealed that disgust sensitivity could not mediate the association between state disgust and the mean activation of the anterior insula cluster (Fig. 3b).

**Fig. 3.**
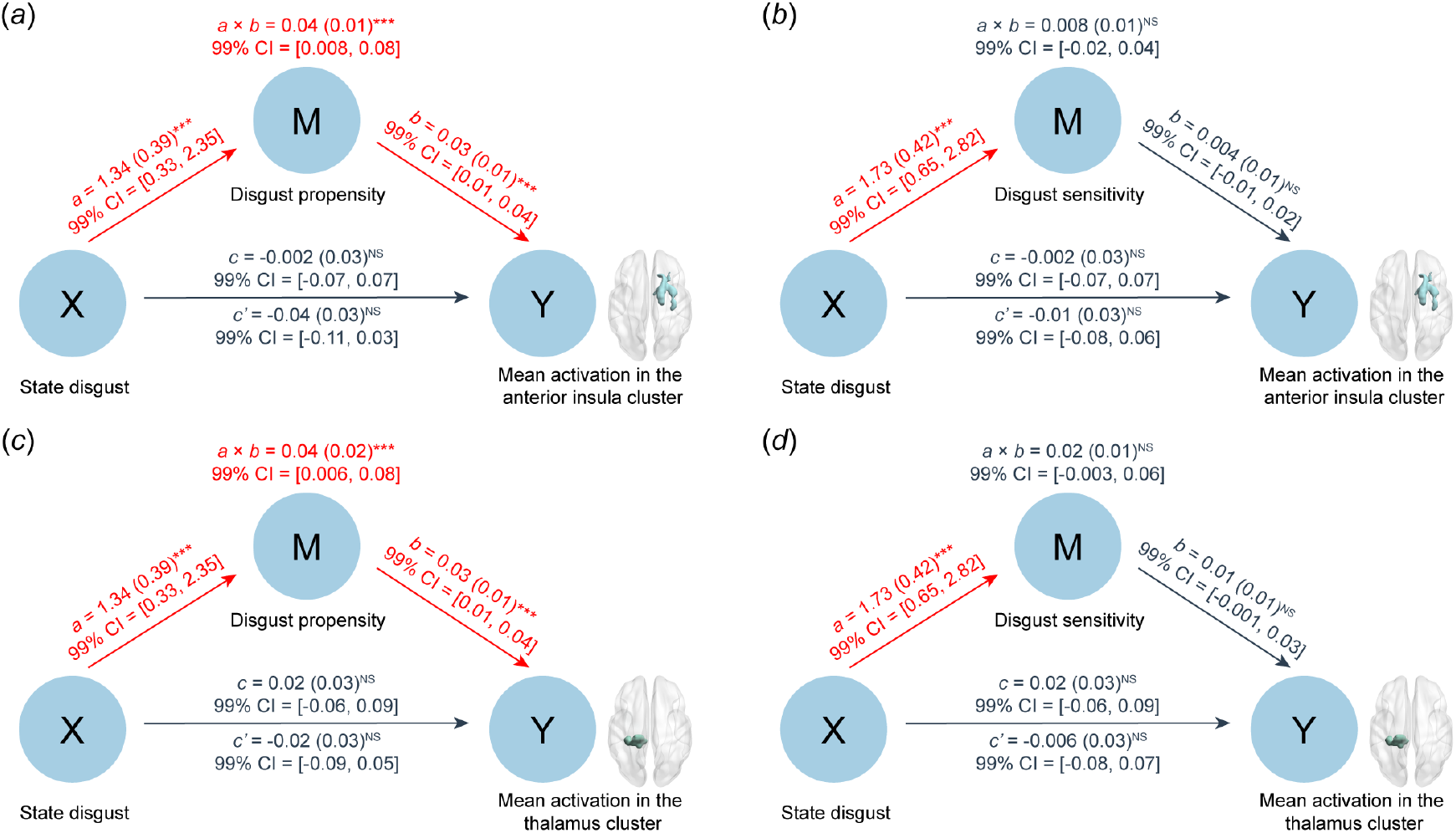
Mediation analyses exploring whether trait disgust mediates the relationship between state disgust and the mean activation in the anterior insula and thalamus clusters. (a) Disgust propensity fully mediates the association between state disgust and the mean activation of the anterior insula cluster. (b) Disgust sensitivity does not mediate the effect of state disgust on the mean activation of the anterior insula cluster. (c) Disgust propensity fully mediates the association between state disgust and the mean activation of the thalamus cluster. (d) Disgust sensitivity fails to mediate the effect of state disgust on the mean activation of the thalamus cluster. \*\*\**P* < 0.001, NS not significant.

We continued to examine whether disgust propensity (*M*) could explain the relationship between state disgust (*X*) and the mean activation of the thalamus cluster (*Y*). According to Fig. 3c, the results showed that state disgust had a positive association with disgust propensity (path a; β = 1.34, SE = 0.39, 99% CI = [0.33, 2.35], *P* < 0.001), disgust propensity was positively correlated with the mean activation of the thalamus cluster (path b; β = 0.03, SE = 0.01, 99% CI = [0.01, 0.04], *P* < 0.001) and there was a significant mediation effect (path a×b; β = 0.04, SE = 0.02, 99% CI = [0.006, 0.08], *P* < 0.001). Moreover, the total effect (path c; β = 0.02, SE = 0.03, 99% CI = [-0.06, 0.09], *P* = 0.53) and the non-mediated effect (path c’; β = −0.02, SE = 0.03, 99% CI = [-0.09, 0.05], *P* = 0.48) were not significant. These results indicate that disgust propensity plays a full mediation role in the effect of state disgust on the mean activation of the thalamus cluster. We next constructed another mediation model with disgust sensitivity as *M*, state disgust as *X*, and the mean activation of the thalamus cluster as *Y*. Nevertheless, the results showed that disgust sensitivity failed to mediate the effect of state disgust on the mean activation of the thalamus cluster (Fig. 3d).

Notably, incorporating the mean activation in each cluster from the regression results after including sex as a covariate in the mediation model (i.e., Table S1) did not affect the pattern of results (Fig. S4).

### (d) Consensus connectivity maps of MACM and RSFC profiles

The consensus connectivity map of the right anterior insula from the regression analysis was determined based on the connectivity profiles of the MACM and RSFC analyses. This analysis yielded a consensus connectivity network of the right anterior insula during both task and resting states, including bilateral anterior and mid- and posterior insular cortex, putamen, caudate, thalamus, and right midcingulate cortex. Moreover, it also contained motor-related regions such as the supplementary motor area, primary motor cortex (e.g., precentral gyrus), and parietal motor regions (e.g., inferior parietal lobule, postcentral gyrus, and supramarginal gyrus) (Fig. 4a, Fig. S5, and Table S4).

**Fig. 4.**
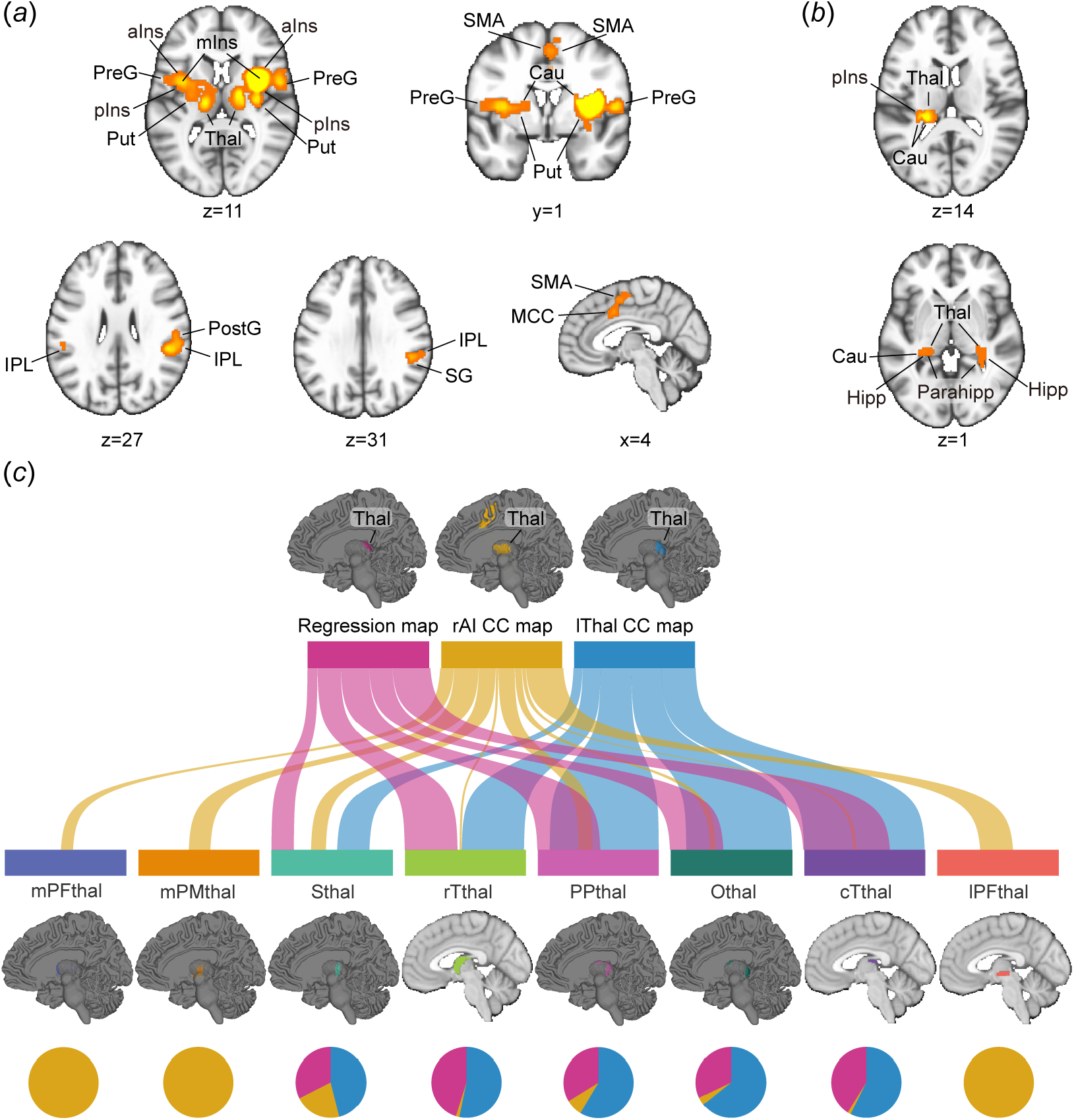
Consensus connectivity networks for the anterior insula and thalamus from the regression analysis. (a) Results of the consensus connectivity map for the right anterior insula. (b) Results of the consensus connectivity map for the left thalamus. (c) River plots showing spatial similarity (cosine similarity) between respective maps (i.e., regression map, Fig. 2; the consensus connectivity map of the right anterior insula, Fig. 4a; the consensus connectivity map of the left thalamus, Fig. 4b) and eight thalamus subregions. Ribbons are normalized by the max cosine similarity across all eight thalamus subregions. Ribbon locations in relation to the boxes are arbitrary. Pie charts show the relative contributions of each map to the thalamus subregion (i.e., percentage of voxels with the highest cosine similarity for each map). Abbreviations: aIns, anterior insula; Cau, caudate; cTthal, caudal temporal thalamus; Hipp, hippocampus; IPL, inferior parietal lobule; lPFthal, lateral prefrontal thalamus; lThal CC map, left thalamus consensus connectivity map; MCC, midcingulate cortex; mIns, middle insula; mPFthal, medial prefrontal thalamus; mPMthal, pre-motor thalamus; Othal, occipital thalamus; Parahipp, parahippocampal gyrus; pIns, posterior insula; PostG, Postcentral Gyrus; PPthal, posterior parietal thalamus; PreG, Precentral Gyrus; Put, putamen; rAI CC map, right anterior insula consensus connectivity map; rTthal, rostral temporal thalamus; SG, supramarginal gyrus; SMA, supplementary motor area; Sthal, sensory thalamus; Thal, thalamus.

Identifying the consensus connectivity map of the left thalamus from the regression analysis showed convergence at the bilateral thalamus, hippocampus, parahippocampal gyrus, left caudate, and left posterior insula (Fig. 4b, Fig. S6, and Table S5).

The following spatial similarity analyses between eight thalamus subregions and the regression map (i.e., Fig. 2) as well as the consensus connectivity maps (i.e., Fig. 4a, b) revealed that the medial prefrontal thalamus, pre-motor thalamus, and lateral prefrontal thalamus uniquely contributed to the consensus connectivity map of the right anterior insula; the sensory thalamus, rostral temporal thalamus, posterior parietal thalamus, occipital thalamus, and caudal temporal thalamus contained significant voxels across all three maps, whereas the latter four thalamus subregions were more strongly involved in the regression map and the consensus connectivity map of the left thalamus (Fig. 4c).

## 4. Discussion

The current study examined whether individual differences in disgust propensity and disgust sensitivity modulate neural responses to disgust-evoking stimuli, using statistically rigorous whole-brain regression analyses in a large, well-powered sample. To account for variations in state disgust, we constructed multiple contrast images reflecting different levels of evoked disgust (e.g., high > neutral, moderate > neutral), which were then entered into separate regression models alongside trait-level measures, thereby allowing us to better isolate their respective modulatory effects on disgust-related neural circuitry. The regression analyses revealed a neural dissociation between the two trait dimensions, with higher levels of disgust propensity – but not variations in disgust sensitivity – being associated with higher disgust-related brain activity in the right insular cortex and adjacent dorsal striatum as well as the left thalamus, caudate, and adjacent (para-)hippocampal formation. Further support for the critical role of these brain systems and the differentiation between the disgust traits was provided by mediation analyses showing that disgust propensity, but not disgust sensitivity, played a full mediation role in the effect of subjective disgust experience (state disgust) on the activation of the anterior insula and thalamic clusters, both of which were identified in the whole-brain regression analyses. Additional network-level results indicated that the two identified regions in the insular cortex and thalamus communicate with distinguishable yet interacting brain systems, with the insular systems engaging with the dorsal striatum, midcingulate cortex, motor-related frontal and parietal regions as well as the thalamus and the thalamic cluster interacting with the posterior insula, (para-)hippocampal formation as well as the caudate. Together, these findings underscore that individual differences in disgust propensity, but not disgust sensitivity, modulate neurofunctional reactivity towards disgust stimuli and mediate the association between neural reactivity in these regions and subjective disgust experience.

The regression analysis revealed that activation in the right anterior insula cluster (including the anterior, mid-, and posterior insula, caudate, and putamen) and the left thalamus cluster (encompassing the thalamus, hippocampus, parahippocampal gyrus, and caudate) showed positive correlations with disgust propensity scores while viewing high disgust stimuli (high disgust > neutral). Among these regions, the insula has been repeatedly defined as a core region in disgust processing. For example, evidence from lesion models [50-54], intracerebral recording studies (e.g., see [55]), and neuroimaging studies in healthy populations with multimodal disgust induction paradigms (e.g., visual, gustatory, olfactory, and imagination [6,9,56-59]) converges to support its role in disgust processing, even though it is a functionally heterogeneous region [35,60-63]. Similarly, basal ganglia regions such as the dorsal striatum, including the caudate and putamen, have been extensively associated with the neural processing of disgust, both in lesion [51,64,65] and multimodal (recall, visual, and gustatory, etc. [25,58,66]) contexts. Although less frequently reported as key disgust-related systems compared insula and basal ganglia, recent meta-analyses have confirmed the role of the parahippocampal gyrus in the experience of disgust [6,9]. The thalamus has not received attention in previous studies on disgust, yet recent studies demonstrate an involvement of the thalamus in anxious arousal [67] and in mediating both subjective experience and autonomous physiological reactivity during affective arousal [34]. Finally, the hippocampus showed a higher reactivity as a function of disgust propensity and, together with the parahippocampal gyrus, may reflect a link between the current emotional experience and memory processing, including valence- and arousal-facilitated mnemonic encoding or an integration of the current affective experience with previous experiences (e.g., see [68-70]). Notably, all these regions observed in the regression analyses have been demonstrated to be associated with and predictive of subjective disgust experience [5]. Overall, these findings may reflect that individual differences in trait disgust propensity modulate neurofunctional reactivity to disgust-inducing stimuli in systems related to the affective experience of disgust and arousal, their interactions with memory, as well as the hard-wired autonomous and motor-preparation responses supporting defensive avoidance reactions.

Further network-level analyses were utilized to determine the networks that the anterior insula and thalamus systems communicate with using a combination of task-based and intrinsic connectivity indices. The consensus connectivity map of the right anterior insula was coupled not only with regions observed in the main regression results (i.e., anterior, mid-, and posterior insular cortex, as well as putamen and caudate), but also with regions such as the thalamus, midcingulate cortex, and a set of motor-related regions (e.g., the supplementary motor area, primary motor cortex, inferior parietal lobule, postcentral gyrus, and supramarginal gyrus). The spatial similarity analyses enabled us to dissect the thalamus in a more nuanced way, revealing that the thalamus in the consensus connectivity map of the right anterior insula primarily comprised the following subparts: medial prefrontal thalamus, lateral prefrontal thalamus, pre-motor thalamus, and sensory thalamus. Functional interactions between these regions may suggest the existence of defensive-avoidance neural circuit that are modulated by disgust propensity. The insula and midcingulate cortex present core nodes of the salience network and may reflect the high salience of disgust-related processes [9], potentially engaging the cortical motor-related regions and subcortical pre-motor thalamus, and, via the basal ganglia and thalamus pathway, initiating goal-directed defensive-avoidance behavior (e.g., withdrawal and inhibition) [53,71-73]. The consensus connectivity map of the left thalamus mirrored the regions identified in the core regression analysis (i.e., thalamus, hippocampus, parahippocampal gyrus, and caudate), and additionally encompassed the posterior insula. Interestingly, spatial similarity analysis revealed that the thalamic subregions in the consensus connectivity map of the left thalamus differed substantially from those observed in the consensus connectivity map of the right anterior insula, but closely matched those identified in the regression analysis (e.g., sensory thalamus, posterior parietal thalamus, occipital thalamus, and rostral and caudal temporal thalamus - the latter four being largely unique to the regression map and the consensus connectivity map of the left thalamus), suggesting strong convergence between trait-linked activation and its connectivity architecture. The above regions observed in the consensus connectivity map of the left thalamus may indicate that the two identified systems centered around the insula and thalamus serve distinct subfunctions in disgust processing, but also converge on the sensory thalamus and posterior insula as relay stations vital for sensory processing [74,75]. Thalamic activity varies with arousal intensities [76] and may initiate autonomic and defensive responses towards potential threats [5,34]; the hippocampus and parahippocampal gyrus may support the integration of the momentary experience with personal and contextual memories related to disgust [68-70,77]; and the engagement of the caudate and posterior insula likely reflects their roles in constructing subjective disgust experience.

The regression analysis incorporating disgust propensity as a covariate for the contrast of high disgust > moderate disgust revealed a set of regions largely overlapping with those observed in the contrast of high disgust > neutral (e.g., the right anterior and left thalamus clusters), while moreover extending to visual and frontal areas. The regression analysis for the moderate disgust > neutral contrast showed that disgust propensity significantly modulated activity in frontal and motor-related regions. Of note, these results are suggestive and should be interpreted cautiously, given the relatively lenient threshold applied (i.e., voxel-level uncorrected *P* < 0.001). These findings also underline the importance of incorporating individual differences in subjective (state) disgust responses, as standard categorizations of stimuli (e.g., ‘high disgust’) may not reflect how strongly each participant actually experiences them [33,34,37,78,79]. Ignoring such variability may blur the mapping between subjective affect and brain activity, and limit our understanding of how trait disgust shapes the emotional neural circuit.

In contrast, no significant effects between disgust sensitivity and disgust-associated brain activity were observed, although on the behavioral level both disgust propensity (*r* = 0.28, *P* < 0.001) and disgust sensitivity (*r* = 0.33, *P* < 0.001) were positively correlated with subjective disgust experience, indicating that behavioral data alone cannot fully distinguish their contributions. This dissociation between disgust propensity and disgust sensitivity at the brain level was further supported by the mediation results showing that it was disgust propensity, rather than disgust sensitivity, that fully mediated the relationship between state disgust and brain responses during disgust-related affective processing, highlighting the added specificity provided by neural measures. Complementary evidence for this dissociation comes from a previous study capitalizing on reverse inference, which demonstrated that the neural signature of subjective disgust experience mediated the influence of the general negative signature on state disgust ratings, but not vice versa [5]. Notably, this study was conducted using the same data as the present work. The differences between disgust sensitivity and disgust propensity observed at the neural level resonate with previous results indicating that the two disgust traits differ in their associations with experience of actual disgust, general negative emotional sensitivity, disgust-related avoidance, and anxiety disorder symptoms [10,11,15,21-24,80].

The findings of this study may also contribute to a better understanding of psychiatric disorders characterized by altered disgust reactivity and disgust-related learning, such as obsessive-compulsive disorder (OCD) [81,82]. On the behavioral level, trait disgust propensity may serve as a psychological marker for monitoring or predicting treatment response in OCD. For instance, Olatunji et al. demonstrated that the reduction in disgust propensity over time mediated improvement in OCD symptoms, even when improvement in negative affect was taken into account [22]. On the neural level, some of the brain regions identified in the present study may represent candidate neural targets for interventions aimed at modulating pathological disgust responses. For example, enhanced activation of the insula cortex and dorsal striatum during processing and generalization of disgust stimuli has been reported in OCD [7,83,84].

Although the present study employed whole-brain regression analyses with stringent threshold corrections and further incorporated mediation analyses to provide tentative mechanistic insights, these methods are inherently correlational in nature and do not permit definitive causal inference. To more robustly establish causal links between trait disgust and neural responses, future research may benefit from noninvasive brain stimulation techniques (e.g., transcranial magnetic stimulation), longitudinal tracking of trait and neural dynamics, or randomized intervention designs targeting trait-level disgust modulation.

In conclusion, previous studies have shown inconsistent results with respect to neural variations related to trait disgust propensity and disgust sensitivity. As the first study to examine the modulatory role of trait disgust on neural responses across varying levels of induced disgust, the current work demonstrates that disgust propensity, rather than disgust sensitivity, robustly modulates neural responses to disgust-inducing stimuli in regions consistently implicated in this emotion. These findings also shed light on the newly established association between heightened disgust propensity and intensified experience of state disgust, by identifying the neural circuit underlying this effect - encompassing the insula, dorsal striatum, thalamus, parahippocampal gyrus, and hippocampus - a network of regions previously linked to defensive and avoidance-related responses to highly aversive stimuli. Furthermore, the mediation and network-level results offer converging support for the above pattern, providing additional insight into how trait disgust shapes affective neural mechanisms. Ultimately, these findings may inform the development of targeted interventions for disgust-related mental disorders.

## References

1. Rozin P., Fallon A.E. 1987 A perspective on disgust. Psychol. Rev. 94, 23–41. (doi:10.1037/0033-295X.94.1.23)

2. Rozin P., Haidt J., Fincher K. 2009 From oral to moral. Science 323, 1179–1180. (doi:10.1126/science.1170492)

3. Tybur J.M., Lieberman D., Kurzban R., DeScioli P. 2013 Disgust: evolved function and structure. Psychol. Rev. 120, 65–84. (doi:10.1037/a0030778)

4. Chapman H.A., Kim D.A., Susskind J.M., Anderson A.K. 2009 In bad taste: evidence for the oral origins of moral disgust. Science 323, 1222–1226. (doi:10.1126/science.1165565)

5. Gan X., Zhou F., Xu T., Liu X., Zhang R., Zheng Z., Yang X., Zhou X., Yu F., Li J., et al. 2024 A neurofunctional signature of subjective disgust generalizes to oral distaste and socio-moral contexts. Nat. Hum. Behav. 8, 1383–1402. (doi:10.1038/s41562-024-01868-x)

6. Gan X., Zhou X., Li J., Jiao G., Jiang X., Biswal B., Yao S., Klugah-Brown B., Becker B. 2022 Common and distinct neurofunctional representations of core and social disgust in the brain: coordinate-based and network meta-analyses. Neurosci. Biobehav. Rev. 135, 104553. (doi:10.1016/j.neubiorev.2022.104553)

7. Vicario C.M., Rafal R.D., Martino D., Avenanti A. 2017 Core, social and moral disgust are bounded: a review on behavioral and neural bases of repugnance in clinical disorders. Neurosci. Biobehav. Rev. 80, 185–200. (doi:10.1016/j.neubiorev.2017.05.008)

8. Jones D. 2007 The depths of disgust. Nature 447, 768–771. (doi:10.1038/447768a)

9. Gan X., Zhang R., Zheng Z., Wang L., Yang X., Klugah-Brown B., Xu T., Qiu N., Kendrick K.M., Mathiak K., et al. 2024 Does unfairness evoke anger or disgust? A quantitative neurofunctional dissection based on 25 years of neuroimaging. bioRxiv, 2024.2010.2017.618853. (doi:10.1101/2024.10.17.618853)

10. van Overveld W.J.M., de Jong P.J., Peters M.L., Cavanagh K., Davey G.C.L. 2006 Disgust propensity and disgust sensitivity: separate constructs that are differentially related to specific fears. Pers. Individ. Differ. 41, 1241–1252. (doi:10.1016/j.paid.2006.04.021)

11. van Overveld M., Jong P.J.d., Peters M.L. 2010 The disgust propensity and sensitivity scale - revised: its predictive value for avoidance behavior. Pers. Individ. Differ. 49, 706–711. (doi:10.1016/j.paid.2010.06.008)

12. Calder A.J., Beaver J.D., Davis M.H., Van Ditzhuijzen J., Keane J., Lawrence A.D. 2007 Disgust sensitivity predicts the insula and pallidal response to pictures of disgusting foods. Eur. J. Neurosci. 25, 3422–3428. (doi:10.1111/j.1460-9568.2007.05604.x)

13. de Jong P.J., Muris P. 2002 Spider phobia: interaction of disgust and perceived likelihood of involuntary physical contact. J. Anxiety Disord. 16, 51–65. (doi:10.1016/S0887-6185(01)00089-5)

14. Deacon B., Olatunji B.O. 2007 Specificity of disgust sensitivity in the prediction of behavioral avoidance in contamination fear. Behav. Res. Ther. 45, 2110–2120. (doi:10.1016/j.brat.2007.03.008)

15. Goetz A.R., Lee H.-J., Cougle J.R., Turkel J.E. 2013 Disgust propensity and sensitivity: differential relationships with obsessive-compulsive symptoms and behavioral approach task performance. Journal of Obsessive-Compulsive and Related Disorders 2, 412–419. (doi:10.1016/j.jocrd.2013.07.006)

16. Olatunji B.O., Armstrong T., Elwood L. 2017 Is disgust proneness associated with anxiety and related disorders? A qualitative review and meta-analysis of group comparison and correlational studies. Perspect. Psychol. Sci. 12, 613–648. (doi:10.1177/1745691616688879)

17. Moretz M.W., McKay D. 2008 Disgust sensitivity as a predictor of obsessive-compulsive contamination symptoms and associated cognitions. J. Anxiety Disord. 22, 707–715. (doi:10.1016/j.janxdis.2007.07.004)

18. Olatunji B.O., Kim J. 2024 Examining reciprocal relations between disgust proneness and OCD symptoms: a four-wave longitudinal study. J. Behav. Ther. Exp. Psychiatry 82, 101907. (doi:10.1016/j.jbtep.2023.101907)

19. van Overveld M., de Jong P.J., Peters M.L., Schouten E. 2011 The Disgust Scale-R: a valid and reliable index to investigate separate disgust domains? Pers. Individ. Differ. 51, 325–330. (doi:10.1016/j.paid.2011.03.023)

20. Aharoni R., Hertz M.M. 2012 Disgust sensitivity and anorexia nervosa. Eur. Eat. Disorders Rev. 20, 106–110. (doi:10.1002/erv.1124)

21. Olatunji B.O., Cisler J., McKay D., Phillips M.L. 2010 Is disgust associated with psychopathology? Emerging research in the anxiety disorders. Psychiatry Res. 175, 1–10. (doi:10.1016/j.psychres.2009.04.007)

22. Olatunji B.O., Tart C.D., Ciesielski B.G., McGrath P.B., Smits J.A. 2011 Specificity of disgust vulnerability in the distinction and treatment of OCD. J. Psychiatr. Res. 45, 1236–1242. (doi:10.1016/j.jpsychires.2011.01.018)

23. Olatunji B.O., Wolitzky-Taylor K.B., Ciesielski B.G., Armstrong T., Etzel E.N., David B. 2009 Fear and disgust processing during repeated exposure to threat-relevant stimuli in spider phobia. Behav. Res. Ther. 47, 671–679. (doi:10.1016/j.brat.2009.04.012)

24. Davey G.C.L. 2011 Disgust: the disease-avoidance emotion and its dysfunctions. Phil. Trans. R. Soc. B 366, 3453–3465. (doi:10.1098/rstb.2011.0039)

25. Mataix-Cols D., An S.K., Lawrence N.S., Caseras X., Speckens A., Giampietro V., Brammer M.J., Phillips M.L. 2008 Individual differences in disgust sensitivity modulate neural responses to aversive/disgusting stimuli. Eur. J. Neurosci. 27, 3050–3058. (doi:10.1111/j.1460-9568.2008.06311.x)

26. Stark R., Schienle A., Sarlo M., Palomba D., Walter B., Vaitl D. 2005 Influences of disgust sensitivity on hemodynamic responses towards a disgust-inducing film clip. Int. J. Psychophysiol. 57, 61–67. (doi:10.1016/j.ijpsycho.2005.01.010)

27. Schienle A., Schäfer A., Stark R., Walter B., Vaitl D. 2005 Relationship between disgust sensitivity, trait anxiety and brain activity during disgust induction. Neuropsychobiology 51, 86–92. (doi:10.1159/000084165)

28. Caseras X., Mataix-Cols D., An S.K., Lawrence N.S., Speckens A., Giampietro V., Brammer M.J., Phillips M.L. 2007 Sex differences in neural responses to disgusting visual stimuli: implications for disgust-related psychiatric disorders. Biol. Psychiatry 62, 464–471. (doi:10.1016/j.biopsych.2006.10.030)

29. Borg C., de Jong P.J., Renken R.J., Georgiadis J.R. 2012 Disgust trait modulates frontal-posterior coupling as a function of disgust domain. Soc. Cogn. Affect. Neurosci. 8, 351–358. (doi:10.1093/scan/nss006)

30. Watkins T.J., Di Iorio C.R., Olatunji B.O., Benningfield M.M., Blackford J.U., Dietrich M.S., Bhatia M., Theiss J.D., Salomon R.M., Niswender K., et al. 2016 Disgust proneness and associated neural substrates in obesity. Soc. Cogn. Affect. Neurosci. 11, 458–465. (doi:10.1093/scan/nsv129)

31. Schäfer A., Leutgeb V., Reishofer G., Ebner F., Schienle A. 2009 Propensity and sensitivity measures of fear and disgust are differentially related to emotion-specific brain activation. Neurosci. Lett. 465, 262–266. (doi:10.1016/j.neulet.2009.09.030)

32. Pujol J., Blanco-Hinojo L., Coronas R., Esteba-Castillo S., Rigla M., Martínez-Vilavella G., Deus J., Novell R., Caixàs A. 2018 Mapping the sequence of brain events in response to disgusting food. Hum. Brain Mapp. 39, 369–380. (doi:10.1002/hbm.23848)

33. Stark R., Zimmermann M., Kagerer S., Schienle A., Walter B., Weygandt M., Vaitl D. 2007 Hemodynamic brain correlates of disgust and fear ratings. Neuroimage 37, 663–673. (doi:10.1016/j.neuroimage.2007.05.005)

34. Zhang R., Gan X., Xu T., Yu F., Wang L., Song X., Jiao G., Liu X., Zhou F., Becker B. 2025 A neurofunctional signature of affective arousal generalizes across valence domains and distinguishes subjective experience from autonomic reactivity. Nat. Commun. 6492. (doi:10.1038/s41467-025-61706-0)

35. Ferraro S., Klugah-Brown B., Tench C.R., Bazinet V., Bore M.C., Nigri A., Demichelis G., Bruzzone M.G., Palermo S., Zhao W., et al. 2022 The central autonomic system revisited -Convergent evidence for a regulatory role of the insular and midcingulate cortex from neuroimaging meta-analyses. Neurosci. Biobehav. Rev. 142, 104915. (doi:10.1016/j.neubiorev.2022.104915)

36. Lang P., Bradley M., Cuthbert B. 2008 International Affective Picture System (IAPS): affective ratings of pictures and instruction manual, Technical Report A-8. University of Florida, Gainesville.

37. Čeko M., Kragel P.A., Woo C.-W., López-Solà M., Wager T.D. 2022 Common and stimulus-type-specific brain representations of negative affect. Nat. Neurosci. 25, 760–770. (doi:10.1038/s41593-022-01082-w)

38. Haberkamp A., Glombiewski J.A., Schmidt F., Barke A. 2017 The DIsgust-RelaTed-Images (DIRTI) database: validation of a novel standardized set of disgust pictures. Behav. Res. Ther. 89, 86–94. (doi:10.1016/j.brat.2016.11.010)

39. Haidt J., McCauley C., Rozin P. 1994 Individual differences in sensitivity to disgust: a scale sampling seven domains of disgust elicitors. Pers. Individ. Differ. 16, 701–713. (doi:10.1016/0191-8869(94)90212-7)

40. Schienle A., Walter B., Stark R., Vaitl D. 2002 Ein Fragebogen zur Erfassung der Ekelempfindlichkeit (FEE). Z. Klin. Psychol. Psychother. 31, 110–120. (doi:10.1026/0084-5345.31.2.110)

41. Marchewka A., Żurawski Ł., Jednoróg K., Grabowska A. 2014 The Nencki Affective Picture System (NAPS): introduction to a novel, standardized, wide-range, high-quality, realistic picture database. Beh. Res. Meth. 46, 596–610. (doi:10.3758/s13428-013-0379-1)

42. Friston K.J., Holmes A.P., Worsley K.J., Poline J.-P., Frith C.D., Frackowiak R.S.J. 1994 Statistical parametric maps in functional imaging: a general linear approach. Hum. Brain Mapp. 2, 189–210. (doi:10.1002/hbm.460020402)

43. Robinson J.L., Laird A.R., Glahn D.C., Lovallo W.R., Fox P.T. 2010 Metaanalytic connectivity modeling: delineating the functional connectivity of the human amygdala. Hum. Brain Mapp. 31, 173–184. (doi:10.1002/hbm.20854)

44. Langner R., Rottschy C., Laird A.R., Fox P.T., Eickhoff S.B. 2014 Meta-analytic connectivity modeling revisited: controlling for activation base rates. Neuroimage 99, 559–570. (doi:10.1016/j.neuroimage.2014.06.007)

45. Nichols T., Brett M., Andersson J., Wager T., Poline J.-B. 2005 Valid conjunction inference with the minimum statistic. Neuroimage 25, 653–660. (doi:10.1016/j.neuroimage.2004.12.005)

46. Bore M.C., Liu X., Gan X., Wang L., Xu T., Ferraro S., Li L., Zhou B., Zhang J., Vatansever D., et al. 2024 Distinct neurofunctional alterations during motivational and hedonic processing of natural and monetary rewards in depression - a neuroimaging meta-analysis. Psychol. Med. 54, 639–651. (doi:10.1017/s0033291723003410)

47. Reimann G.M., Hoseini A., Koçak M., Beste M., Küppers V., Rosenzweig I., Elmenhorst D., Pires G.N., Laird A.R., Fox P.T., et al. 2025 Distinct Convergent Brain Alterations in Sleep Disorders and Sleep Deprivation: a Meta-Analysis. JAMA Psychiatry 82, 681–691. (doi:10.1001/jamapsychiatry.2025.0488)

48. Fan L., Li H., Zhuo J., Zhang Y., Wang J., Chen L., Yang Z., Chu C., Xie S., Laird A.R., et al. 2016 The human brainnetome atlas: a new brain atlas based on connectional architecture. Cereb. Cortex 26, 3508–3526. (doi:10.1093/cercor/bhw157)

49. Zhang X., Qing P., Liu Q., Liu C., Liu L., Gan X., Fu K., Lan C., Zhou X., Kendrick K.M., et al. 2025 Neural patterns of social pain in the brain-wide representations across social contexts. Advanced Science, 2413795. (doi:10.1002/advs.202413795)

50. Adolphs R., Tranel D., Damasio A.R. 2003 Dissociable neural systems for recognizing emotions. Brain Cogn. 52, 61–69. (doi:10.1016/S0278-2626(03)00009-5)

51. Calder A.J., Keane J., Manes F., Antoun N., Young A.W. 2000 Impaired recognition and experience of disgust following brain injury. Nat. Neurosci. 3, 1077–1078. (doi:10.1038/80586)

52. Cantone M., Lanza G., Bella R., Pennisi G., Santalucia P., Bramanti P., Pennisi M. 2019 Fear and disgust: case report of two uncommon emotional disturbances evoked by visual disperceptions after a right temporal-insular stroke. BMC Neurol. 19, 193. (doi:10.1186/s12883-019-1417-0)

53. Holtmann O., Bruchmann M., Mönig C., Schwindt W., Melzer N., Miltner W.H.R., Straube T. 2020 Lateralized deficits of disgust processing after insula-basal ganglia damage. Front. Psychol. 11, 1429. (doi:10.3389/fpsyg.2020.01429)

54. Kipps C.M., Duggins A.J., McCusker E.A., Calder A.J. 2007 Disgust and happiness recognition correlate with anteroventral insula and amygdala volume respectively in preclinical Huntington’s disease. J. Cogn. Neurosci. 19, 1206–1217. (doi:10.1162/jocn.2007.19.7.1206)

55. Krolak-Salmon P., Hénaff M.-A., Isnard J., Tallon-Baudry C., Guénot M., Vighetto A., Bertrand O., Mauguière F. 2003 An attention modulated response to disgust in human ventral anterior insula. Ann. Neurol. 53, 446–453. (doi:10.1002/ana.10502)

56. Heining M., Young A., Ioannou G., Andrew C., Brammer M., Gray J., Phillips M. 2004 Disgusting smells activate human anterior insula and ventral striatum. Ann. N. Y. Acad. Sci. 1000, 380–384. (doi:10.1196/annals.1280.035)

57. Wicker B., Keysers C., Plailly J., Royet J.-P., Gallese V., Rizzolatti G. 2003 Both of us disgusted in my insula: the common neural basis of seeing and feeling disgust. Neuron 40, 655–664. (doi:10.1016/S0896-6273(03)00679-2)

58. Jabbi M., Bastiaansen J., Keysers C. 2008 A common anterior insula representation of disgust observation, experience and imagination shows divergent functional connectivity pathways. PLoS ONE 3, e2939. (doi:10.1371/journal.pone.0002939)

59. Corradi-DellAcqua C., Tusche A., Vuilleumier P., Singer T. 2016 Cross-modal representations of first hand and vicarious pain, disgust and fairness in insular and cingulate cortex. Nat. Commun. 7, 10904. (doi:10.1038/ncomms10904)

60. Li J., Xu L., Zheng X., Fu M., Zhou F., Xu X., Ma X., Li K., Kendrick K.M., Becker B. 2019 Common and dissociable contributions of alexithymia and autism to domain-specific interoceptive dysregulations: a dimensional neuroimaging approach. Psychother. Psychosom. 88, 187–189. (doi:10.1159/000495122)

61. Molnar-Szakacs I., Uddin L.Q. 2022 Anterior insula as a gatekeeper of executive control. Neurosci. Biobehav. Rev. 139, 104736. (doi:10.1016/j.neubiorev.2022.104736)

62. Uddin L.Q. 2015 Salience processing and insular cortical function and dysfunction. Nat. Rev. Neurosci. 16, 55–61. (doi:10.1038/nrn3857)

63. Zhou F., Li J., Zhao W., Xu L., Zheng X., Fu M., Yao S., Kendrick K.M., Wager T.D., Becker B. 2020 Empathic pain evoked by sensory and emotional-communicative cues share common and process-specific neural representations. eLife 9, e56929. (doi:10.7554/eLife.56929)

64. Paulmann S., Pell M.D., Kotz S.A. 2009 Comparative processing of emotional prosody and semantics following basal ganglia infarcts: ERP evidence of selective impairments for disgust and fear. Brain Res. 1295, 159–169. (doi:10.1016/j.brainres.2009.07.102)

65. Gray J.M., Young A.W., Barker W.A., Curtis A., Gibson D. 1997 Impaired recognition of disgust in Huntington’s disease gene carriers. Brain 120, 2029–2038. (doi:10.1093/brain/120.11.2029)

66. Fitzgerald D.A., Posse S., Moore G.J., Tancer M.E., Nathan P.J., Phan K.L. 2004 Neural correlates of internally-generated disgust via autobiographical recall: a functional magnetic resonance imaging investigation. Neurosci. Lett. 370, 91–96. (doi:10.1016/j.neulet.2004.08.007)

67. Liu X., Jiao G., Zhou F., Kendrick K.M., Yao D., Gong Q., Xiang S., Jia T., Zhang X.-Y., Zhang J., et al. 2024 A neural signature for the subjective experience of threat anticipation under uncertainty. Nat. Commun. 15, 1544. (doi:10.1038/s41467-024-45433-6)

68. Xu T., Zhou X., Jiao G., Zeng Y., Zhao W., Li J., Yu F., Zhou F., Yao S., Becker B. 2022 Angiotensin antagonist inhibits preferential negative memory encoding via decreasing hippocampus activation and its coupling with the amygdala. Biol. Psychiatry Cogn. Neurosci. Neuroimaging 7, 970–978. (doi:10.1016/j.bpsc.2022.05.007)

69. Pacheco-Estefan D., Bouyeure A., Jacob G., Fellner M.-C., Lehongre K., Lambrecq V., Frazzini V., Navarro V., Güntürkün O., Shen L., et al. 2025 Representational dynamics during extinction of fear memories in the human brain. Nat. Hum. Behav. (doi:10.1038/s41562-025-02268-5)

70. Qasim S.E., Mohan U.R., Stein J.M., Jacobs J. 2023 Neuronal activity in the human amygdala and hippocampus enhances emotional memory encoding. Nat. Hum. Behav. 7, 754–764. (doi:10.1038/s41562-022-01502-8)

71. Mair R.G., Francoeur M.J., Krell E.M., Gibson B.M. 2022 Where actions meet outcomes: medial prefrontal cortex, central thalamus, and the basal ganglia. Front. Behav. Neurosci. 16, 928610. (doi:10.3389/fnbeh.2022.928610)

72. Martinez-Garcia R.I., Voelcker B., Zaltsman J.B., Patrick S.L., Stevens T.R., Connors B.W., Cruikshank S.J. 2020 Two dynamically distinct circuits drive inhibition in the sensory thalamus. Nature 583, 813–818. (doi:10.1038/s41586-020-2512-5)

73. Liu X., Lai J., Han C., Zhong H., Huang K., Liu Y., Zhu X., Wei P., Tan L., Xu F., et al. 2025 Neural circuit underlying individual differences in visual escape habituation. Neuron 113, 2344-2357.e2345. (doi:10.1016/j.neuron.2025.04.018)

74. Torrico T.J., Munakomi S. 2023 Neuroanatomy, Thalamus, StatPearls Publishing, Treasure Island (FL).

75. Alonso-Matielo H., Zhang Z., Gambeta E., Huang J., Chen L., de Melo G.O., Dale C.S., Zamponi G.W. 2023 Inhibitory insula-ACC projections modulate affective but not sensory aspects of neuropathic pain. Mol. Brain 16, 64. (doi:10.1186/s13041-023-01052-8)

76. Anders S., Lotze M., Erb M., Grodd W., Birbaumer N. 2004 Brain activity underlying emotional valence and arousal: a response-related fMRI study. Hum. Brain Mapp. 23, 200–209. (doi:10.1002/hbm.20048)

77. Tao D., He Z., Lin Y., Liu C., Tao Q. 2021 Where does fear originate in the brain? A coordinate-based meta-analysis of explicit and implicit fear processing. Neuroimage 227, 117686. (doi:10.1016/j.neuroimage.2020.117686)

78. Zhou F., Zhao W., Qi Z., Geng Y., Yao S., Kendrick K.M., Wager T.D., Becker B. 2021 A distributed fMRI-based signature for the subjective experience of fear. Nat. Commun. 12, 6643. (doi:10.1038/s41467-021-26977-3)

79. Chang L.J., Gianaros P.J., Manuck S.B., Krishnan A., Wager T.D. 2015 A sensitive and specific neural signature for picture-induced negative affect. PLoS Biol. 13, e1002180. (doi:10.1371/journal.pbio.1002180)

80. Cisler J.M., Olatunji B.O., Lohr J.M. 2009 Disgust sensitivity and emotion regulation potentiate the effect of disgust propensity on spider fear, blood-injection-injury fear, and contamination fear. J. Behav. Ther. Exp. Psychiatry 40, 219–229. (doi:10.1016/j.jbtep.2008.10.002)

81. Wang J., Becker B., Wang Y., Ming X., Lei Y., Wikgren J. 2024 Conceptual-level disgust conditioning in contamination-based obsessive-compulsive disorder. Psychophysiology 61, e14637. (doi:10.1111/psyp.14637)

82. Rast C., Woronko S., Jessup S.C., Olatunji B.O. 2023 Treatment of disgust in specific emotional disorders. Bull. Menninger Clin. 87, 5–30. (doi:10.1521/bumc.2023.87.suppA.5)

83. Liu J., Wang J., Song Y., Becker B., Ming X., Lei Y. 2025 Enhanced disgust generalization in obsessive-compulsive disorder is related to insula and putamen hyperactivity. Psychol. Med. 55, e116. (doi:10.1017/s0033291725000728)

84. Shapira N.A., Liu Y., He A.G., Bradley M.M., Lessig M.C., James G.A., Stein D.J., Lang P.J., Goodman W.K. 2003 Brain activation by disgust-inducing pictures in obsessive-compulsive disorder. Biol. Psychiatry 54, 751–756. (doi:10.1016/S0006-3223(03)00003-9)

