## Supplemental Materials for "Disgust propensity, not disgust sensitivity, shapes the reactivity of a subjective disgust circuit in humans"

### Supplementary Methods

#### MRI data acquisition

The MRI data of the first experiment were acquired on a GE MR750 (General Electric Medical System) 3T MRI system. Functional images were obtained with a T2*-weighted echo planar imaging (EPI) sequence with the following parameters: repetition time = 2000 ms, echo time = 30 ms, flip angle = 90°, field of view = 240 × 240 mm, voxel size = 3.75 × 3.75 × 4 mm, resolution = 64 × 64, number of slices = 39, slice thickness = 3 mm, and an inter-slice gap of 1 mm. High-resolution structural images were acquired using a T1-weighted spoiled gradient recall (SPGR) sequence with the following parameters: repetition time = 6 ms, echo time = 2 ms, flip angle = 9°, field of view = 256 × 256 mm, voxel size = 1 × 1 × 1 mm, acquisition matrix = 256 × 256, number of slices = 156, and slice thickness = 1 mm. These structural images were utilized to enhance spatial normalization and to exclude participants with evident brain pathologies.

The MRI data of the second experiment were acquired on a GE MR750 3T MRI system. Functional images were obtained with a T2*-weighted EPI sequence with the following parameters: repetition time = 2000 ms, echo time = 30 ms, flip angle = 90°, field of view = 200 × 200 mm, voxel size = 3.125×3.125×3.8 mm, resolution = 64 × 64, number of slices = 36, slice thickness = 3.8 mm, and no gap. High-resolution structural images were acquired using a T1-weighted SPGR sequence with the following parameters: repetition time = 8 ms, echo time = 3 ms, flip angle = 8°, field of view = 256 × 256 mm, voxel size = 1 × 1 × 1 mm, acquisition matrix = 256 × 256, number of slices = 176, and slice thickness = 1 mm. These structural images were used to improve spatial normalization and to exclude participants with apparent brain pathologies.

#### Statistical rationale of mediation analysis

Mediation analysis aims to investigate whether the association between an independent variable (X) and a dependent variable (Y) can be explained by a third variable, known as the mediator (M). The X-M relation, M-Y relation, and X-Y relation before and after controlling for the M are characterized by paths a, b, c, and c′, respectively. The total effect of X on Y (path c) is the sum of the direct effect (path c′) and the indirect (mediation) effect (the product of the path coefficients of paths a and b, i.e., a × b). Evidence for mediation is established when paths a, b, and a × b are all statistically significant. Moreover, when c′ is not significant, M plays a full mediation role; otherwise, M plays a partial mediation role.

#### Activation likelihood estimation (ALE)

The ALE method estimates the probability of each voxel being activated across multiple independent experiments, taking into account the number of participants in each study. In other words, ALE seeks to determine whether the convergence of activation foci across included experiments occurs at a level greater than chance [1]. Specifically, ALE treats the reported peak foci from single studies as three-dimensional Gaussian probability distributions and provides empirical estimates to address the spatial uncertainty that may arise due to between-subject and between-template variability of the neuroimaging foci [2]. For each Gaussian probability distribution function, the full width at half maxima is weighted by the sample size of the respective experiment, such that experiments with larger samples produce smaller Gaussian distributions, reflecting more precise and reliable estimates of “true” activation [2].

For each included study, modeled activation maps were generated by combining the probabilities of all activation foci for each voxel [2,3]. The union of individual modeled activation maps across experiments was calculated to produce an ALE map, reflecting brain regions (clusters) where activation convergence was observed [2,4]. Consistent with standard practices, the ALE map was thresholded using a cluster-level familywise error (cluster-FWE) correction at $P < 0.05$, with an initial cluster-forming threshold set at the voxel-level of $P < 0.001$ (uncorrected) [5-11]. The $P$ value reflects the proportion of the null distribution of random spatial relation across experiments. In our analyses, $P$ values were generated by 5000 permutations [10,12,13].

**Data acquisition parameters and preprocessing procedures of the OASIS-3 dataset**

Images were acquired on a Siemens TimTrio 3T scanner. Resting-state functional images were obtained using an echo-planner imaging sequence using the following parameters: repetition time = 2200 ms, echo time = 27 ms, flip angle = 90°, voxel size = 4.0 × 4.0 × 4.0 $mm^3$, 33 slices, and slice thickness = 4 mm. Preprocessing was conducted using Data Processing Assistant for Resting-State fMRI (DPARSF, v.5.0_201001) (http://rfmri.org/DPARSF). In brief, the preprocessing pipeline included the following steps: (1) discarding the first five volumes to improve stability of the magnetic field, (2) slice-timing correction to account for temporal differences in slice acquisition, (3) realignment to correct for head motion, (4) normalization of functional images to the standard MNI space, (5) grey matter, white matter, cerebrospinal fluid, and head motion regression, (6) detrending, (7) filtering (0.01 - 0.1 Hz), and (8) spatial smoothing with a 6-mm full-width at half-maximum Gaussian kernel.

**Supplementary Results**

**Regression results for the condition of high disgust > moderate disgust**

Applying the same threshold (i.e., cluster-level FWE $P < 0.05$ correction and voxel-level $P < 0.001$) to the condition of high disgust > moderate disgust did not yield any results. Given this, we reported the regression results at the voxel-level $P < 0.001$ to observe some trend. The results showed that activation to high disgust stimuli (high disgust > moderate disgust) was significantly modulated by participants' disgust propensity scores within the bilateral anterior, mid- and posterior insular cortex, bilateral occipital visual cortex, right putamen, left thalamus, left hippocampus, left parahippocampal gyrus, left caudate, and left inferior/middle frontal gyrus (Table S2 and Fig. S2).

**Regression results for the condition of moderate disgust > neutral**

Applying the same threshold (i.e., cluster-level FWE $P < 0.05$ correction and voxel-level $P < 0.001$) to the condition of moderate disgust > neutral failed to yield any results. We thus relax the threshold to voxel-level $P < 0.001$. The results showed that activation to moderate disgust stimuli (moderate disgust > neutral) was significantly modulated by participants' disgust propensity scores in the right precentral gyrus, right postcentral gyrus, bilateral paracentral lobule, and right middle/superior frontal gyrus (Table S3 and Fig. S3).

Table S1. Brain areas where activation to viewing high disgust stimuli (high disgust > neutral) was significantly associated with participants' scores on the disgust propensity subscale (further including sex as a covariate in the regression analysis)

| Regions | Side | MNI peak coordinates (x, y, z) | | | *t*-value | Cluster Size | Cluster-FWE |
|---|---|---|---|---|---|---|---|
| Anterior Insula, Middle Insula, Posterior Insula, Caudate, Putamen | R | 34 | 4 | 14 | 4.58 | 546 | 0.009 |
| Thalamus, Hippocampus, Caudate, Parahippocampal Gyrus | L | -24 | -32 | 10 | 4.71 | 477 | 0.016 |

Note: voxel-level uncorrected *P* < 0.001, combined with cluster-level FWE correction. Abbreviations: L, left; R, right.

Table S2. Brain areas where activation to viewing high disgust stimuli (high disgust > moderate disgust) was significantly associated with participants' scores on the disgust propensity subscale

| Regions | Side | MNI peak coordinates (x, y, z) | | | *t*-value | Cluster Size |
|---|---|---|---|---|---|---|
| Anterior Insula, Middle Insula, Posterior Insula, Putamen | R | 34 | 4 | 14 | 5.67 | 222 |
| Thalamus | L | -14 | -26 | 14 | 4.44 | 112 |
| Hippocampus, Parahippocampal Gyrus, Caudate | L | -34 | -32 | -2 | 4.18 | 144 |
| Inferior Occipital Gyrus, Lingual Gyrus, Middle Occipital Gyrus | L | -18 | -96 | -6 | 4.00 | 151 |
| Anterior Insula, Middle Insula, Posterior Insula | L | -36 | 0 | 14 | 3.84 | 51 |
| Middle Frontal Gyrus, Inferior Frontal Gyrus | L | -32 | 32 | 16 | 3.72 | 57 |
| Anterior Insula, Middle Insula | L | -30 | 12 | 14 | 3.70 | 37 |
| Lingual Gyrus, Inferior Occipital Gyrus, Middle Occipital Gyrus | R | 20 | -96 | -4 | 3.51 | 118 |

Note: voxel-level uncorrected *P* < 0.001. Abbreviations: L, left; R, right.

Table S3. Brain areas where activation to viewing moderate disgust stimuli (moderate disgust > neutral) was significantly associated with participants' scores on the disgust propensity subscale

| Regions | Side | MNI peak coordinates (x, y, z) | | | *t*-value | Cluster Size |
|---|---|---|---|---|---|---|
| Precentral Gyrus | R | 46 | 0 | 32 | 3.88 | 55 |
| Precentral Gyrus, Postcentral Gyrus | R | 44 | -18 | 68 | 3.78 | 45 |
| Superior Frontal Gyrus, Middle Frontal Gyrus | R | 26 | 58 | -20 | 3.77 | 61 |
| Paracentral Lobule | R/L | 4 | -30 | 60 | 3.62 | 68 |

Note: voxel-level uncorrected $P < 0.001$. Abbreviations: L, left; R, right.

Table S4. Conjunction analysis of MACM and RSFC results for the right anterior insula from regression analysis

| Regions | Side | MNI peak coordinates (x, y, z) | | | Cluster Size |
|---|---|---|---|---|---|
| Anterior Insula, Middle Insula, Posterior Insula, Putamen, Caudate, Thalamus, Precentral Gyrus | R | 34 | 4 | 12 | 1394 |
| Thalamus, Anterior Insula, Middle Insula, Posterior Insula, Putamen, Caudate, Precentral Gyrus | L | -14 | -16 | 6 | 775 |
| Inferior Parietal Lobule, Supramarginal Gyrus, Postcentral Gyrus | R | 48 | -34 | 26 | 233 |
| Inferior Parietal Lobule | L | -54 | -30 | 24 | 13 |
| Midcingulate Cortex, Supplementary Motor Area | R/L | 8 | 12 | 38 | 297 |

Abbreviations: L, left; R, right.

Table S5. Conjunction analysis of MACM and RSFC results for the left thalamus from regression analysis

| Regions | Side | MNI peak coordinates (x, y, z) | | | Cluster Size |
|---|---|---|---|---|---|
| Thalamus, Hippocampus, Posterior Insula, Caudate, Parahippocampal Gyrus | L | -22 | -32 | 8 | 384 |
| Thalamus, Hippocampus, Parahippocampal Gyrus | R | 28 | -32 | 2 | 56 |

Abbreviations: L, left; R, right.

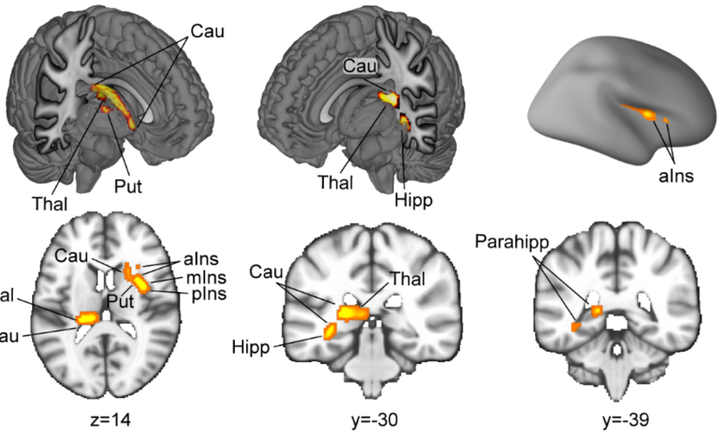


Fig. S1 Positive association between disgust propensity scores and brain activation while viewing high disgust stimuli (high disgust > neutral), with sex as a covariate. Note: activated voxel group shown at a statistical threshold of voxel-level *P* < 0.001 and cluster-level *P* < 0.05. Abbreviations: aIns, anterior insula; Cau, caudate; Hipp, hippocampus; mIns, middle insula; Parahipp, parahippocampal gyrus; pIns, posterior insula; Put, putamen; Thal, thalamus.

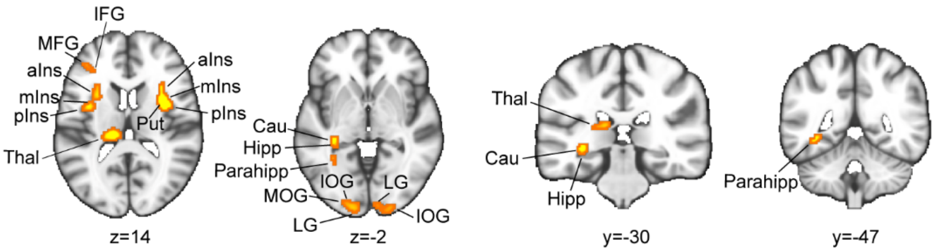


Fig. S2 Positive association between disgust propensity scores and brain activation while viewing high disgust stimuli (high disgust > moderate disgust). Note: activated voxel group shown at a statistical threshold of voxel-level *P* < 0.001. Abbreviations: aIns, anterior insula; Cau, caudate; Hipp, hippocampus; IFG, inferior frontal gyrus; IOG, inferior occipital gyrus; LG, lingual gyrus; MFG, middle frontal gyrus; mIns, middle insula; MOG, middle occipital gyrus; Parahipp, parahippocampal gyrus; pIns, posterior insula; Put, putamen; Thal, thalamus.

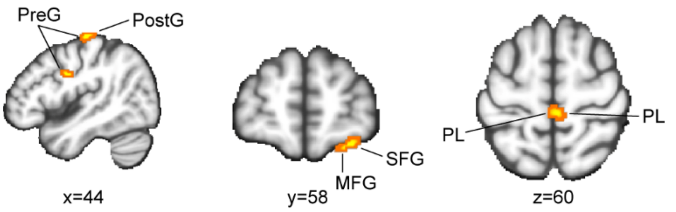


Fig. S3 Positive association between disgust propensity scores and brain activation while viewing moderate disgust stimuli (moderate disgust > neutral). Note: activated voxel group shown at a statistical threshold of voxel-level *P* < 0.001. Abbreviations: MFG, middle frontal gyrus; PL, paracentral lobule; PostG, postcentral gyrus; PreG, precentral gyrus; SPG, superior frontal gyrus.

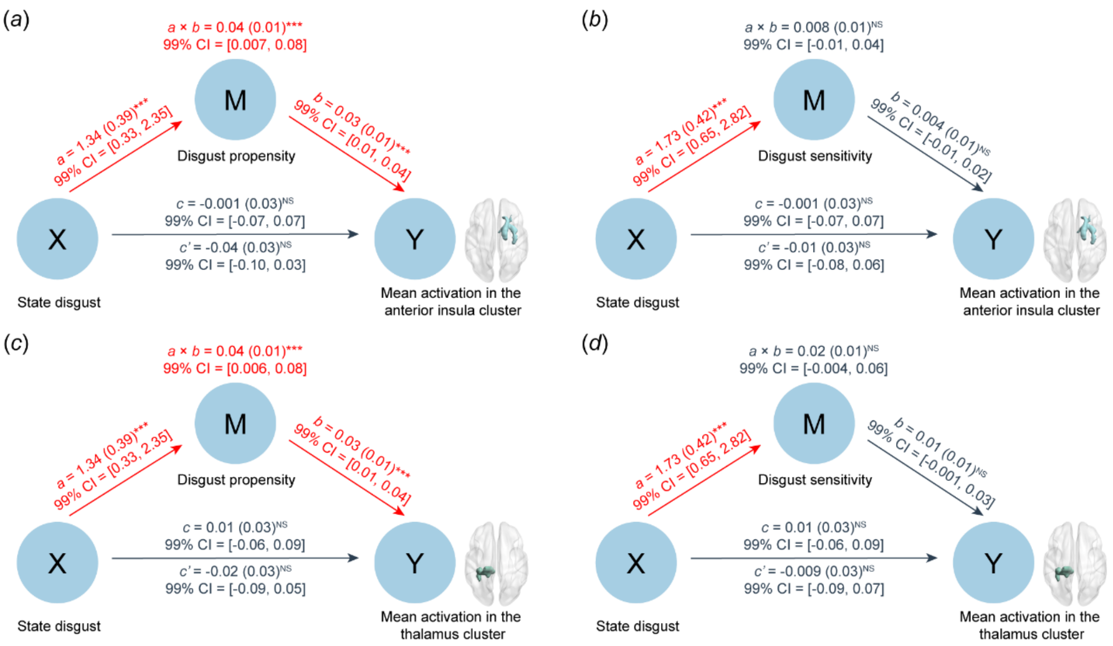


Fig. S4 Mediation analyses exploring whether trait disgust mediates the relationship between state disgust and the mean activation (from regression results that included sex as a covariate) in the anterior insula and thalamus clusters. (a) Disgust propensity fully mediates the association between state disgust and the mean activation of the anterior insula cluster. (b) Disgust sensitivity does not mediate the effect of state disgust on the mean activation of the anterior insula cluster. (c) Disgust propensity fully mediates the association between state disgust and the mean activation of the thalamus cluster. (d) Disgust sensitivity fails to mediate the effect of state disgust on the mean activation of the thalamus cluster. ****P* < 0.001, NS not significant.

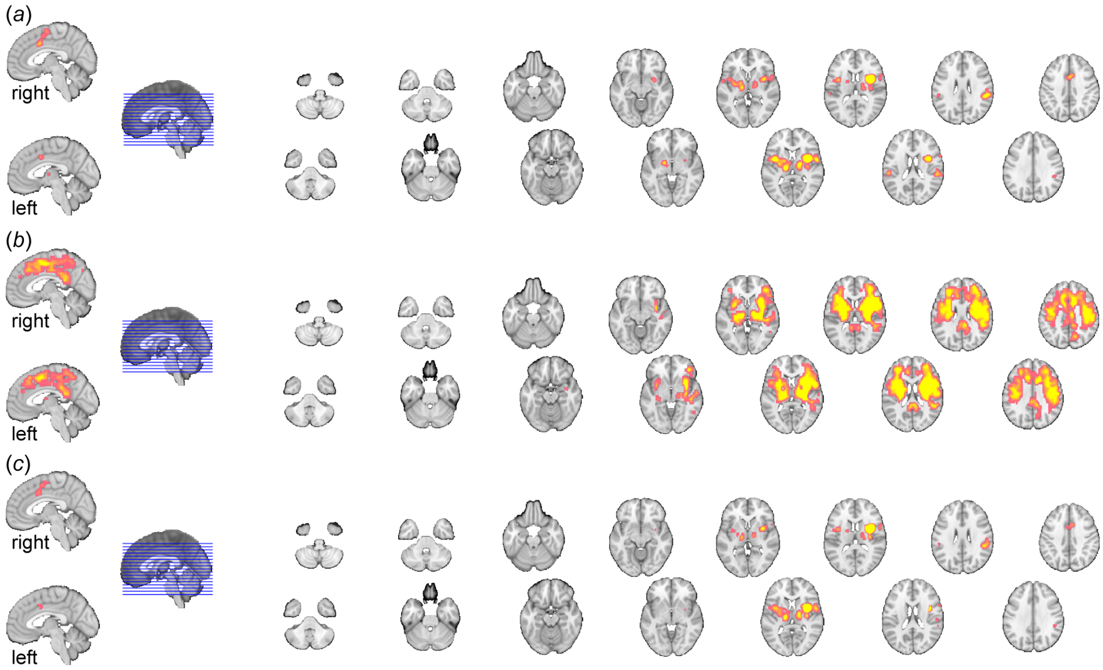


Fig. S5 Connectivity profiles of the right anterior insula from regression analysis. (a) Task-based connectivity map (i.e., MACM). (b) Task-free connectivity map (i.e., RSFC). (c) Consensus connectivity map (i.e., MACM ∩ RSFC, where ∩ indicates conjunction).

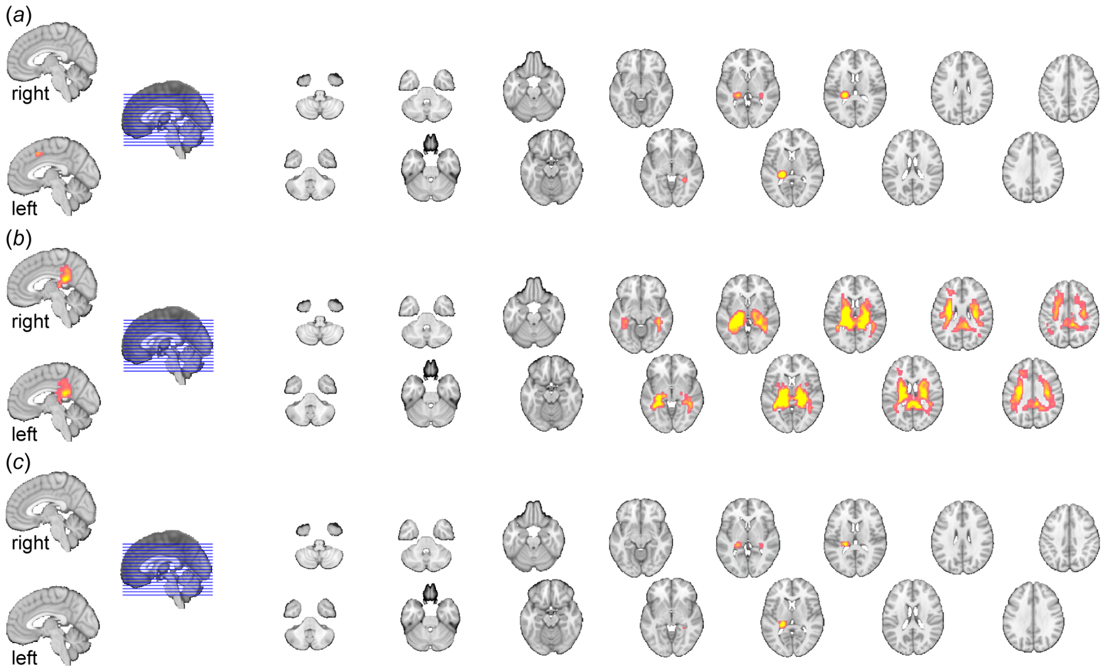


Fig. S6 Connectivity profiles of the left thalamus from regression analysis. (a) Task-based connectivity map (i.e., MACM). (b) Task-free connectivity map (i.e., RSFC). (c) Consensus connectivity map (i.e., MACM ∩ RSFC, where ∩ indicates conjunction).

### References


1. Laird A.R., Fox P.M., Price C.J., Glahn D.C., Uecker A.M., Lancaster J.L., Turkeltaub P.E., Kochunov P., Fox P.T. 2005 ALE meta-analysis: controlling the false discovery rate and performing statistical contrasts. *Hum. Brain Mapp.* **25**, 155-164. (doi:10.1002/hbm.20136)
2. Eickhoff S.B., Laird A.R., Grefkes C., Wang L.E., Zilles K., Fox P.T. 2009 Coordinate-based activation likelihood estimation meta-analysis of neuroimaging data: a random-effects approach based on empirical estimates of spatial uncertainty. *Hum. Brain Mapp.* **30**, 2907-2926. (doi:10.1002/hbm.20718)
3. Turkeltaub P.E., Eickhoff S.B., Laird A.R., Fox M., Wiener M., Fox P. 2012 Minimizing within-experiment and within-group effects in activation likelihood estimation meta-analyses. *Hum. Brain Mapp.* **33**, 1-13. (doi:10.1002/hbm.21186)
4. Eickhoff S.B., Bzdok D., Laird A.R., Kurth F., Fox P.T. 2012 Activation likelihood estimation meta-analysis revisited. *Neuroimage* **59**, 2349-2361. (doi:10.1016/j.neuroimage.2011.09.017)
5. Eickhoff S.B., Laird A.R., Fox P.M., Lancaster J.L., Fox P.T. 2017 Implementation errors in the GingerALE software: description and recommendations. *Hum. Brain Mapp.* **38**, 7-11. (doi:10.1002/hbm.23342)
6. Eickhoff S.B., Nichols T.E., Laird A.R., Hoffstaedter F., Amunts K., Fox P.T., Bzdok D., Eickhoff C.R. 2016 Behavior, sensitivity, and power of activation likelihood estimation characterized by massive empirical simulation. *Neuroimage* **137**, 70-85. (doi:10.1016/j.neuroimage.2016.04.072)
7. Müller V.I., Cieslik E.C., Laird A.R., Fox P.T., Radua J., Mataix-Cols D., Tench C.R., Yarkoni T., Nichols T.E., Turkeltaub P.E., et al. 2018 Ten simple rules for neuroimaging meta-analysis. *Neurosci. Biobehav. Rev.* **84**, 151-161. (doi:10.1016/j.neubiorev.2017.11.012)
8. Klugah-Brown B., Di X., Zweerings J., Mathiak K., Becker B., Biswal B. 2020 Common and separable neural alterations in substance use disorders: A coordinate-based meta-analyses of functional neuroimaging studies in humans. *Hum. Brain Mapp.* **41**, 4459-4477. (doi:10.1002/hbm.25085)
9. Klugah-Brown B., Zhou X., Pradhan B.K., Zweerings J., Mathiak K., Biswal B., Becker B. 2021 Common neurofunctional dysregulations characterize obsessive-compulsive, substance use, and gaming disorders - an activation likelihood meta-analysis of functional imaging studies. *Addict. Biol.* **26**, e12997. (doi:10.1111/adb.12997)
10. Gan X., Zhou X., Li J., Jiao G., Jiang X., Biswal B., Yao S., Klugah-Brown B., Becker B. 2022 Common and distinct neurofunctional representations of core and social disgust in the brain: coordinate-based and network meta-analyses. *Neurosci. Biobehav. Rev.* **135**, 104553. (doi:10.1016/j.neubiorev.2022.104553)
11. Ferraro S., Klugah-Brown B., Tench C.R., Bazinet V., Bore M.C., Nigri A., Demichelis G., Bruzzone M.G., Palermo S., Zhao W., et al. 2022 The central autonomic system revisited - Convergent evidence for a regulatory role of the insular and midcingulate cortex from neuroimaging meta-analyses. *Neurosci. Biobehav. Rev.* **142**, 104915. (doi:10.1016/j.neubiorev.2022.104915)
12. Laird A.R., Robinson J.L., McMillan K.M., Tordesillas-Gutiérrez D., Moran S.T., Gonzales S.M., Ray K.L., Franklin C., Glahn D.C., Fox P.T., et al. 2010 Comparison of the disparity between Talairach and MNI coordinates in functional neuroimaging data: validation of the Lancaster transform. *Neuroimage* **51**, 677-683. (doi:10.1016/j.neuroimage.2010.02.048)
13. Tao D., He Z., Lin Y., Liu C., Tao Q. 2021 Where does fear originate in the brain? A coordinate-based meta-analysis of explicit and implicit fear processing. *Neuroimage* **227**, 117686.

(doi:10.1016/j.neuroimage.2020.117686)
